# Attention drives visual processing and audiovisual integration during multimodal communication

**DOI:** 10.1101/2023.05.11.540320

**Authors:** Noor Seijdel, Jan-Mathijs Schoffelen, Peter Hagoort, Linda Drijvers

## Abstract

During communication in real-life settings, our brain often needs to integrate auditory and visual information, and at the same time actively focus on the relevant sources of information, while ignoring interference from irrelevant events. The interaction between integration and attention processes remains poorly understood. Here, we use rapid invisible frequency tagging (RIFT) and magnetoencephalography (MEG) to investigate how attention affects auditory and visual information processing and integration, during multimodal communication. We presented human participants (male and female) with videos of an actress uttering action verbs (auditory; tagged at 58 Hz) accompanied by two movie clips of hand gestures on both sides of fixation (attended stimulus tagged at 65 Hz; unattended stimulus tagged at 63 Hz). Integration difficulty was manipulated by a lower-order auditory factor (clear/degraded speech) and a higher-order visual semantic factor (matching/mismatching gesture). We observed an enhanced neural response to the attended visual information during degraded speech compared to clear speech. For the unattended information, the neural response to mismatching gestures was enhanced compared to matching gestures. Furthermore, signal power at the intermodulation frequencies of the frequency tags, indexing non-linear signal interactions, was enhanced in left frontotemporal and frontal regions. Focusing on LIFG, this enhancement was specific for the attended information, for those trials that benefitted from integration with a matching gesture. Higher power at this intermodulation frequency was related to faster reaction times. Together, our results suggest that attention modulates the strength and speed of audiovisual processing and interaction, depending on the congruence and quality of the sensory input.

## Introduction

In daily conversations, our brains are bombarded with sensory input from various modalities, making it impossible to comprehensively process everything and everyone in our environment. To effectively communicate in real-life settings, we must not only process auditory information, such as speech, and visual information, like mouth movements and co-speech gestures, but also selectively attend to relevant sources of information while ignoring irrelevant ones. The extent to which the integration of audiovisual speech information is automatic, or influenced by diverted attention conditions, is still a topic of debate (for reviews see: Navarra et al., 2010; Koelewijn et al., 2010; Talsma et al., 2010; Macaluso et al., 2016). While some studies have demonstrated that audiovisual integration is a rather unavoidable process, even when the relevant stimuli are outside the focus of attention (Foxe et al., 2000; Driver 1996; Bertelson et al., 2000; Vroomen et al., 2001a, 2001b), others have shown that audiovisual integration is vulnerable to diverted attention conditions or to visually crowded scenarios (Ahmed et al., 2021; Alsius et al., 2005; 2007; 2014; Alsius and Soto-Faraco, 2011, Andersen Tobias et al., 2009; Buchan and Munhall, 2011; 2012; Fairhall and Macaluso, 2009; Fujisaki et al., 2006; Senkowski et al., 2005; Tiippana et al., 2011). Thus, how audiovisual integration and attention interact remains poorly understood.

Recent developments put forward a new technique, Rapid Invisible Frequency Tagging (RIFT), as an important tool to investigate exactly this question. RIFT enables researchers to track both attention to multiple stimuli, and investigate the integration of audiovisual signals (Drijvers et al., 2021; Seijdel et al., 2023; Brickwedde et al., 2022; Minarik et al., 2022; Pan et al., 2021; Duecker et al., 2021; Marshall et al., 2021; Zhigalov et al., 2019;2021; Zhigalov & Jensen, 2020; Ferrante et al., 2023). This technique, in which visual stimuli are periodically modulated at high (>50 Hz), stimulus specific ‘tagging frequencies’, generates steady-state evoked potentials with strong power at the tagged frequencies (Norcia et al., 2015; Vialatte et al., 2010). Frequency tagging has been shown to be a flexible technique to investigate the tracking of attention to multiple different stimuli, with a functional relationship between the amplitude of the SSVEP and the deployment of attention (Toffanin et al. 2009), reflecting the benefit of spatial attention on perceptual processing (Zhigalov et al., 2019). Frequency tagging is interesting in the context of studying audiovisual integration, to investigate whether and how auditory and visual input interact in the brain. Tagging simultaneously presented auditory (using e.g. amplitude modulation) and visual stimuli at different frequencies may lead to non-linear signal interactions indexed by a change in signal power at so-called intermodulation frequencies. For example, using RIFT and magnetoencephalography (MEG), Drijvers et al. (2021) identified an intermodulation frequency at 7 Hz (f_visual_ − f_auditory_) as a result of the interaction between a visual frequency-tagged signal (gesture; 68 Hz) and an auditory frequency-tagged signal (speech; 61 Hz).

In the present study, we investigated how attention affects the processing of auditory and visual information, as well as their integration, during multimodal stimulus presentation. Specifically, we used RIFT and MEG to measure neural activity in response to videos of an actress uttering action verbs (auditory) accompanied with visual gestures on both sides of fixation. We manipulated integration difficulty by varying a lower-order auditory factor (clear/degraded speech) and a higher-order visual factor (congruent/incongruent gesture) and tagged the stimuli at different frequencies for the attended and unattended stimuli. We expected power in visual regions to reflect attention towards the visually tagged input. For the auditory input, we expected power in auditory regions reflecting attention to the auditory tagged input. We expected the interaction between the (attended and ignored) visually tagged signals and the auditory tagged signal to result in spectral peaks at the intermodulation frequencies (65-58 and 63-58; 7 Hz and 5 Hz) respectively. Specifically, we expected this peak to be higher for the attended information (7 Hz) and we expected this activity to occur in the left inferior frontal gyrus (LIFG), a region known to be involved in speech-gesture integration.

## Materials and methods

### Participants

Forty participants (20 females, 18-40 years old) took part in the experiment. Data from two participants were excluded after data collection, due to missed exclusion criteria (one participant was too old) and problems with comprehension of the task instructions (one participant always answered using the visual information as leading information). All remaining participants were right-handed and reported corrected-to-normal or normal vision. None of the participants had language, motor or neurological impairment and all reported normal hearing. All participants gave written consent before they participated in the experiment. Participants received monetary compensation or research credits for their participation. The study was approved by the local ethical committee (CMO:2014/288).

### Stimuli

The same stimuli as in Drijvers et al. (2021) were used. Participants were presented with 160 video clips showing an actress uttering a highly-frequent action verb accompanied by a matching or a mismatching iconic gesture. Auditory information could be clear or degraded and visual information (gestures) could be congruent or incongruent. In total, there were four conditions, each consisting of 40 trials: clear speech + matching gesture (CM), clear speech, mismatching gesture (CMM), degraded speech + matching gesture (DM) and degrading speech + mismatching gesture (DMM). In all videos, the actress was standing in front of a neutrally colored curtain, in neutrally colored clothes.

During recording of the videos, all gestures were performed by the actress on the fly (the gestures were not predetermined). Verbs for the mismatching gestures were predefined to allow the actress to utter the action verb and depict the mismatching gesture while the face and lips still matched the speech. Videos were on average 2000 ms long. After 120 ms, the preparation (i.e., the first frame in which the hands of the actress moved) of the gesture started. On average, at 550 ms the meaningful part of the gesture (i.e., the stroke) started, followed by speech onset at 680 ms, and average speech offset at 1435 ms. None of these timings differed between conditions. None of the iconic gestures were prescripted. All audio files were intensity-scaled to 70 dB and denoised using Praat (Boersma & Weenink, 2015), before they were recombined with their corresponding video files using Adobe Premiere Pro. To degrade the audio, files were noise-vocoded using Praat. Noise-vocoding preserves the temporal envelope of the audio signal, but degrades the spectral content (Shannon et al., 1995). Based on previous work (Drijvers & Ozyürek, 2017), we used 6-band noise-vocoding, to ensure participants still were able to understand enough of the auditory features of the speech signal to integrate the visual semantic information from the gesture. For further details and descriptions see Drijvers et al., 2017 and Drijvers et al., 2021.

### Experimental design and statistical analyses

Participants were tested in a dimly-lit magnetically shielded room and seated 70 cm from the projection screen. All stimuli were presented using MATLAB 2016b (Mathworks Inc, Natrick, USA) and the Psychophysics Toolbox, version 3.0.11 (Brainard, 1997; Kleiner, Brainard, & Pelli, 2007). To achieve RIFT, we used a GeForce GTX960 2GB graphics card with a refresh rate of 120 Hz, in combination with a PROPixx DLP LED projector (VPixx Technologies Inc., Saint-Bruno-de-Montarville, Canada), which can achieve a presentation rate up to 1,440 Hz. This high presentation rate is achieved by the projector interpreting the four quadrants and three color channels of the GPU screen buffer as individual smaller, grayscale frames, which it then projects in rapid succession, leading to an increase of a factor 12 (4 quadrants * 3 color channels * 120 Hz = 1,440 Hz). The area of the video that would be frequency-tagged was defined by the rectangle in which all gestures occurred. This was achieved by multiplying the luminance of the pixels within that square with a 65/63 Hz sinusoid (modulation depth = 100%; modulation signal equal to 0.5 at sine wave zero-crossing, in order to preserve the mean luminance of the video), phase-locked across trials. For the auditory stimuli, frequency tagging was achieved by multiplying the amplitude of the signal with a 58 Hz sinusoid, with a modulation depth of 100% (following Drijvers et al 2021; Lamminmäki, Parkkonen, & Hari, 2014).

To manipulate spatial attention, we added an attentional cue (arrow pointing to the left or right presented before video onset) and presented the same visual stimulus twice, with different tagging frequencies left and right of fixation. We used the same video to avoid unwanted effects from different properties of the videos (e.g. differences in salience, movement kinematics). On half of the trials, participants were asked to attend to the video on the left side of fixation, on the other half of the trials participants were asked to attend to the video on the right side of fixation. The attended video was frequency-tagged at 65 Hz, the unattended video at 63 Hz. The area of the videos that would be frequency-tagged was defined by the rectangle in which all gestures occurred (see Drijvers et al. 2021 for full procedure). Participants were asked to attentively watch and listen to the videos. Every trial started with a fixation cross (1000 ms), followed by the attentional cue (1000 ms), the videos (2000 ms), a short delay period (1500 ms), and a 4-alternative forced choice identification task (max 3000 ms, followed by the fixation cross of the next trial as soon as a participant pressed one of the 4 buttons). In the 4-alternative forced choice identification task, participants were presented with four written options, and had to identify which verb they heard in the video by pressing one of 4 buttons on an MEG-compatible button box (Figure 1). These answering options always contained a phonological distractor, a semantic distractor, an unrelated answer, and the correct answer. For example, the correct answer could be “strikken” (to tie), the phonological distractor could be “tikken” (to tick), the semantic distractor, which would fit with the gesture, could be “knopen” (to button), and the unrelated answer could be “zouten” (to salt). This task ensured that participants were attentively watching the videos, and enabled us to check whether the verbs were understood. Participants were instructed not to blink during video presentation. The stimuli were presented in four blocks of 40 trials each. In addition to the normal trials, “attention trials” were included to stimulate and monitor attention (see Figure 1). During these trials, participants performed an orthogonal task using already presented stimuli. In these trials, a change in brightness could occur in the attended video, at different latencies. Participants were asked to detect this change in brightness. The whole experiment lasted ∼30 minutes and participants were allowed to take a self-paced break after every block. All stimuli were presented in a randomized order per participant.

**Figure 1.**
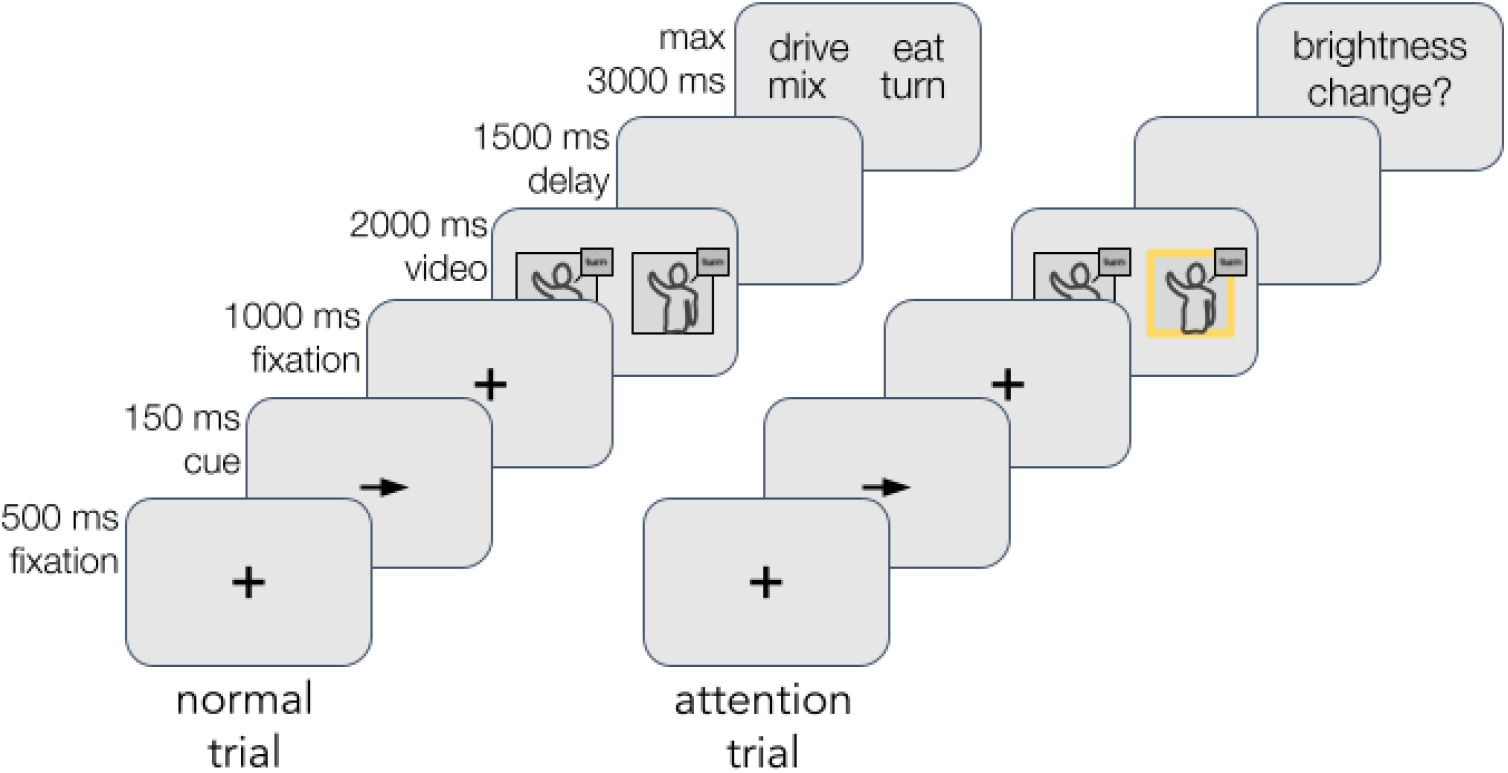
Experimental paradigm. Participants were asked to attend to one of the videos, indicated by a cue. The attended video was frequency-tagged at 65 Hz, the unattended video at 63 Hz. Speech was frequency-tagged at 58 Hz. Participants were asked to attentively watch and listen to the videos. A line drawing of the actress and video setup is shown for illustration purposes. After the video, participants were presented with four written options, and had to identify which verb they heard in the video by pressing one of 4 buttons on an MEG-compatible button box. This task ensured that participants were attentively watching the videos, and to check whether the verbs were understood. Participants were instructed not to blink during the video presentation. In addition to the normal trials, “attention trials” were included in which participants were asked to detect a change in brightness.

#### Data acquisition

Brain activity was measured using MEG, and was recorded throughout the experiment. MEG was acquired using a whole-brain CTF-275 system with axial gradiometers (CTF MEG systems, Coquitlam, Canada). Data were sampled at 1200 Hz after a 300 Hz low-pass filter was applied. Six sensors (MRF66, MLC11, MLC32, and MLO33, MRO33 and MLC61) were permanently disabled due to high noise. Head location was measured using localization coils in both ear canals and on the nasion and was monitored continuously using online head localization software (Stolk et al., 2013). In case of large deviations from the initial head position, we paused the experiment and instructed the subject to move back to the original position. Participants’ eye gaze was recorded by an SR Research Eyelink 1,000 eye tracker for artifact rejection purposes. During the task, participants responded using a Fiber Optic Response Pad placed at their right hand.

After the experiment, T1-weighted anatomical magnetic resonance images (MRI) were acquired in the sagittal orientation (or obtained in case of previous participation in MRI/MEG research) using a 3D MPRAGE sequence with the following parameters: TR/TI/TE = 2300/1100/3 ms, FA = 8º, FOV = 256x225x192 mm and a 1 mm isotropic resolution. Parallel imaging (iPAT = 2) was used to accelerate the acquisition resulting in an acquisition time of 5 min and 21 sec. To align structural MRI to MEG, we placed vitamin E capsules in the external meatus of the ear canals, at the same locations as the localizer coils in the MEG system. These anatomical scans were used for source reconstruction of the MEG signals.

#### Behavioral analysis

Choice accuracy and reaction times (RT) were computed for each condition and each participant. RT analysis was performed on correct responses only. RTs < 100 ms were considered “fast guesses” and removed. Behavioral data were analyzed in Python using the following packages: Statsmodels, Pingouin, SciPy, NumPy, Pandas, (Jones et al., 2001; Vallat, 2018; Oliphant, 2006; Seabold and Perktold, 2010; McKinney, 2011).

#### MEG preprocessing

MEG data were preprocessed and analyzed using the FieldTrip toolbox (Oostenveld et al., 2011) and custom-built MATLAB scripts (2021b). The MEG signal was epoched based on the onset of the video (t = -1 to 3 s). The data were downsampled to a sampling frequency of 400 Hz after applying a notch filter to remove line noise and harmonics (50, 100, 150, 200, 250, 300 and 350 Hz). Bad channels and trials were rejected via a semi-automatic routine before independent-component analysis (Bell et al., 1995; Jung et al., 2001) was applied. Subsequently, components representing eye-related and heart-related artifacts were projected out of the data (on average, 3.7 components were removed per participant). These procedures resulted in rejection of 9.3% of the trials. The number of rejected trials did not differ significantly between conditions.

#### Frequency Tagging - sensor and source

We first evaluated power at the tagging frequencies in visual and auditory sensory areas by calculating power spectra in the stimulus time window (0.5-1.5 s) and the post-stimulus time window (2.0-3.0 s). We chose this post-stimulus time window as a baseline because, contrary to the prestimulus time window, it is not affected by the button press of the 4-alternative forced choice identification task (following the procedure in Drijvers et al., 2021). To facilitate interpretation of the MEG data, we calculated synthetic planar gradients, as planar gradient maxima are known to be located above neural sources that may underlie them (Bastiaansen & Knösche, 2000). For each individual and each condition, we conducted a spectral analysis for all frequencies between 1 and 130 Hz with a step size of 1 Hz. We applied the fast Fourier transform to the planar-transformed time domain data, after tapering with a boxcar window. Afterward, the horizontal and vertical components of the planar gradient were combined by summing. Using the power spectrum during the baseline condition, the percentage increase in power during stimulus presentation was computed. The resulting power per frequency was averaged over participants and visualized. For the auditory tagging we evaluated all temporal sensors, for the visual tagging we evaluated all occipital channels. Then, to investigate whether RIFT can be used to identify intermodulation frequencies as a result of the interaction between visual and auditory tagged signals, we repeated the procedure and evaluated power at the intermodulation frequencies (5 Hz and 7 Hz). Here, we focused on left frontal sensors, as the left frontal cortex is known to be involved in the integration of speech and gesture.

Source analysis was performed using dynamic imaging of coherent sources (DICS; Gross et al., 2001). DICS computes source level power at specified frequencies for a set of predefined locations. For each of these locations a beamformer spatial filter is constructed from the sensor-level cross-spectral density matrix (CSD) and the location’s lead field matrix. We obtained individual lead fields for every participant using the anatomical information from their MRI. First, we spatially co-registered the individual anatomical MRI to sensor space MEG data by identifying the anatomical markers at the nasion and the two ear canals. We then constructed a realistically shaped single-shell volume conduction model on the basis of the segmented MRI for each participant, and divided the brain volume into a 10 mm spaced grid and warped it to a template brain (MNI). To evaluate power spectra in our sensory regions-of-interest (ROIs) we evaluated visual tagging in all occipital channels, and auditory tagging in all temporal channels. At source level, we evaluated visual tagging in occipital cortex, including all occipital regions involved with visual processing based on the The Human Brainnetome Atlas (regions 189-196; 199-201; Fan et al. 2016). Auditory tagging was evaluated in temporal regions A41/42 and A22 (regions 71, 72, 75,76,79, 80).

Next, we zoomed in on the tagging frequencies and identified the sources of the oscillatory activity. After establishing regions that showed enhanced power at the tagging- and intermodulation frequencies, we proceeded to test the effect of the experimental conditions (clear vs. degraded speech; matching vs. mismatching gesture) within these regions-of-interest (ROIs). The ROIs for the auditory and visual tagged signals were defined by taking the grid points that exceeded 80% of the peak power difference value between stimulus and baseline, across all conditions. For these ROIs, power difference values were extracted per condition. Based on previous studies, the ROI for the intermodulation frequencies at 5 and 7 Hz was anatomically defined by taking those grid points that were part of the Left Inferior Frontal Gyrus (LIFG), using the The Human Brainnetome Atlas; (Fan et al. 2016). To evaluate whether power at the intermodulation frequencies in LIFG was increased during the stimulus window compared to the post-stimulus baseline window, 1 sample permutation tests against zero were performed, using 5000 permutations. For each permutation the signs of a random number of entries in the sample were flipped and the difference in means from the null population mean was recomputed. We repeated this until all permutations were evaluated and stored the differences. The p-value was computed by taking the number of times the stored differences were at least as extreme as the original difference, divided by the total number of permutations. In each iteration, all samples were taken into account (resampling was dependent only on the assignment of values to condition groups)

#### Linking neural activity to behavior

Finally, we investigated whether the power at the intermodulation frequency was related to the speed of successful recognition behavior (faster responses). This would indicate that the observed intermodulation frequency in higher-order regions reflects the ease of the integration of audiovisual stimuli.

## Results

In the behavioral task we replicated previous results (see Drijvers, Ozyürek, et al., 2018; Drijvers & Özyürek, 2018; Drijvers, Jensen, Spaak, 2021) and observed that when the speech signal was clear, response accuracy was higher than when speech was degraded (F[1, 37]= 649.82, *p*<.001, partial *η*^2^ = 0.946). Participants performed better when the gesture matched the speech signal compared to when the gesture mismatched the speech signal (F[1, 37]=39.95, *p*<.001, partial *η*^2^ = 0.519). There was a significant interaction between Speech (clear/degraded) and Gesture (matching/mismatching) (F[1, 37]=46.30, *p*<.001, partial *η*^2^ = 0.556). Gestures hindered comprehension when the actress performed a mismatching gesture and speech was degraded (Figure 2).

**Figure 2.**
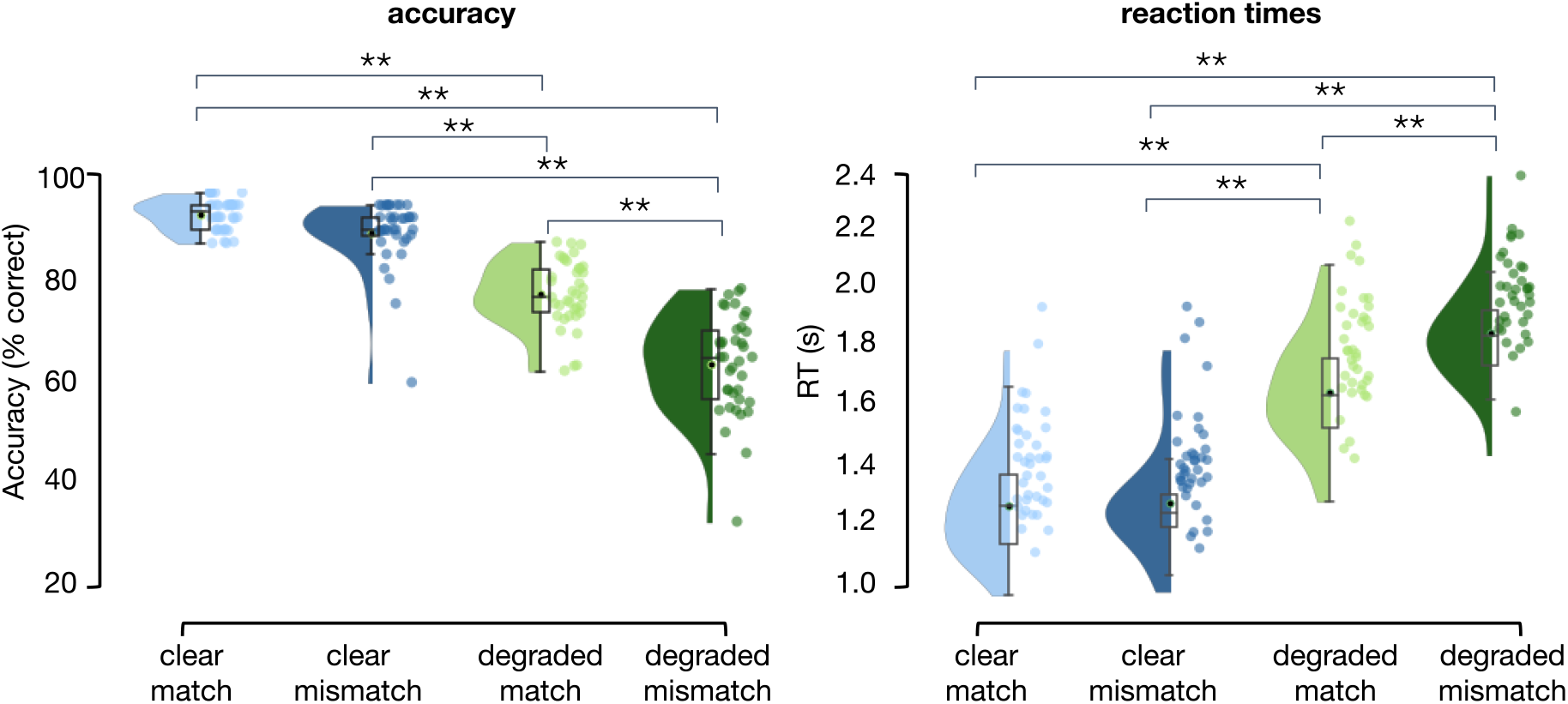
Verb categorization behavior. **A)** Accuracy results per condition. Response accuracy is highest for clear speech conditions, and when a gesture matches the speech signal. **B)** Reaction times per condition. Reaction times are faster in clear speech and when a gesture matches the speech signal.

We observed similar results in the RTs. Participants were faster to identify the verbs when speech was clear, compared to when speech was degraded (F[1, 37] = 568.76, *p* <.001, partial *η*^2^ = 0.939). Participants were also faster to identify the verbs when the gesture matched the speech signal, compared to when the gesture mismatched the speech signal (F[1, 37] = 31.04, *p* < .001, partial *η*^2^ = 0.456). There was a significant interaction between Speech (clear/degraded) and Gesture (matching/mismatching) (F[1, 37] = 47.41, *p* <.001, partial *η*2 = 0.562). Gestures slowed responses when the actress performed a mismatching gesture and speech was degraded.

In sum, these results demonstrate that the presence of a matching or a mismatching gesture modulates speech comprehension. This effect was larger in degraded speech than in clear speech.

### Both visual and auditory frequency tagging produced a clear response that is larger than baseline

Both visual and auditory frequency tagging produced a clear steady-state response that was larger than baseline. A one-sample permutation test against zero with 5000 permutations indicated that for the temporal sensors, spectral power was increased at the auditory tagging frequency, 58 Hz (Figure 3A), *p* < .001. For occipital sensors power was increased at the visual tagging frequencies, 63 and 65 Hz (Figure 3B), *p* < .001 and *p* < .001, respectively. We confirmed these results at the source level, by computing the source spectra to evaluate power at the different frequencies in our regions of interest (based on the The Human Brainnetome Atlas; (Fan et al. 2016). Robust tagging responses were found over auditory cortex (58 Hz; Figure 3C) and visual cortex (65 Hz, 63 Hz; Figure 3D), reflecting the neural resources associated with auditory and visual processing.

**Figure 3.**
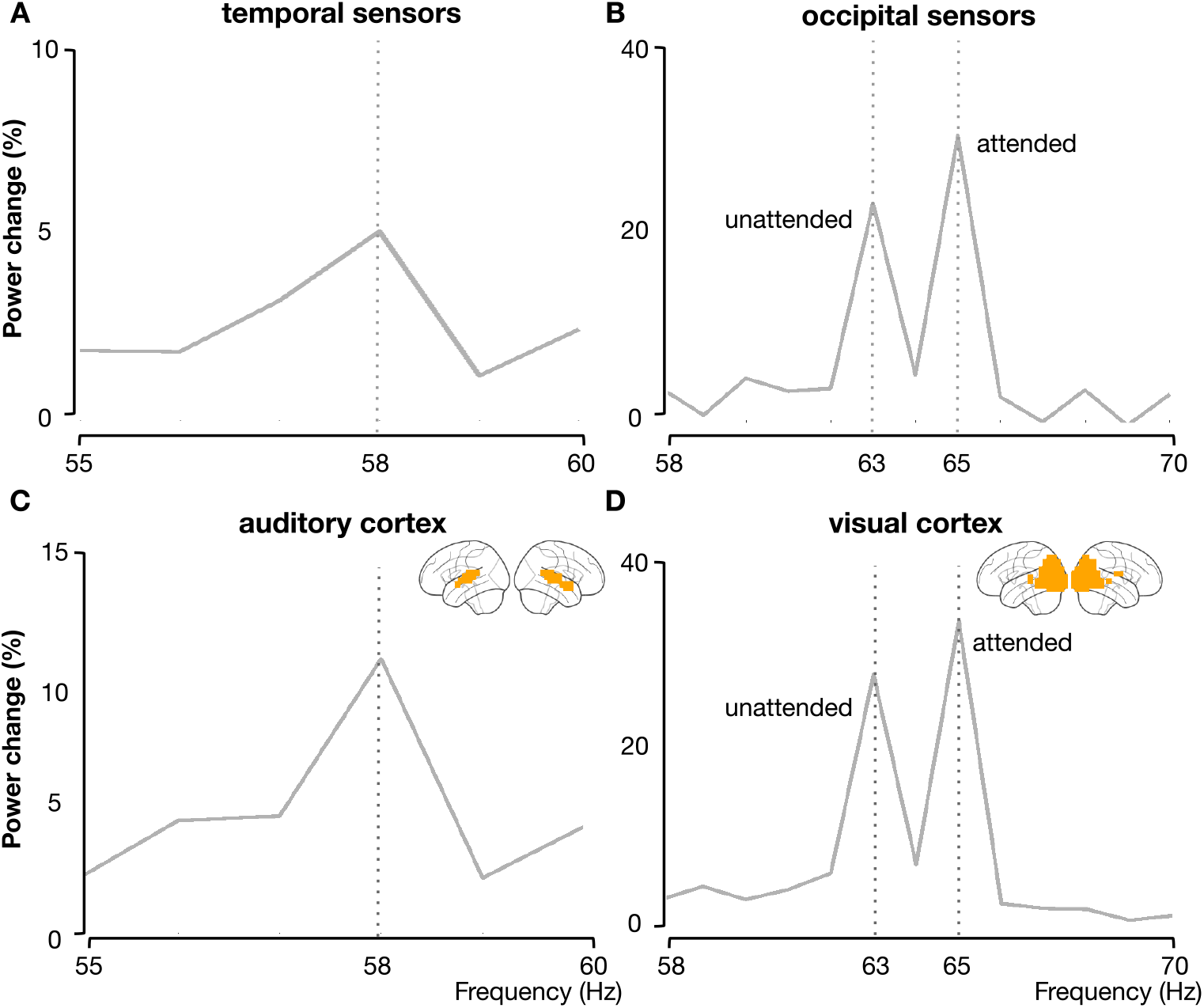
Power at temporal and occipital sensors and corresponding source regions (% increased compared to a post-stimulus baseline) averaged across conditions. **A)** power increase in temporal sensors and in auditory cortex at the tagged frequency of the auditory stimulus (58 Hz) **B)** power increases in occipital sensors are observed at the visual tagging frequencies (63 Hz: unattended; 65 Hz: attended). **C)** power increase in auditory cortex at the tagged frequency of the auditory stimulus (58 Hz). D) power increases in visual cortex observed at the visual tagging frequencies (63 Hz: unattended; 65 Hz: attended).

**Figure 4.**
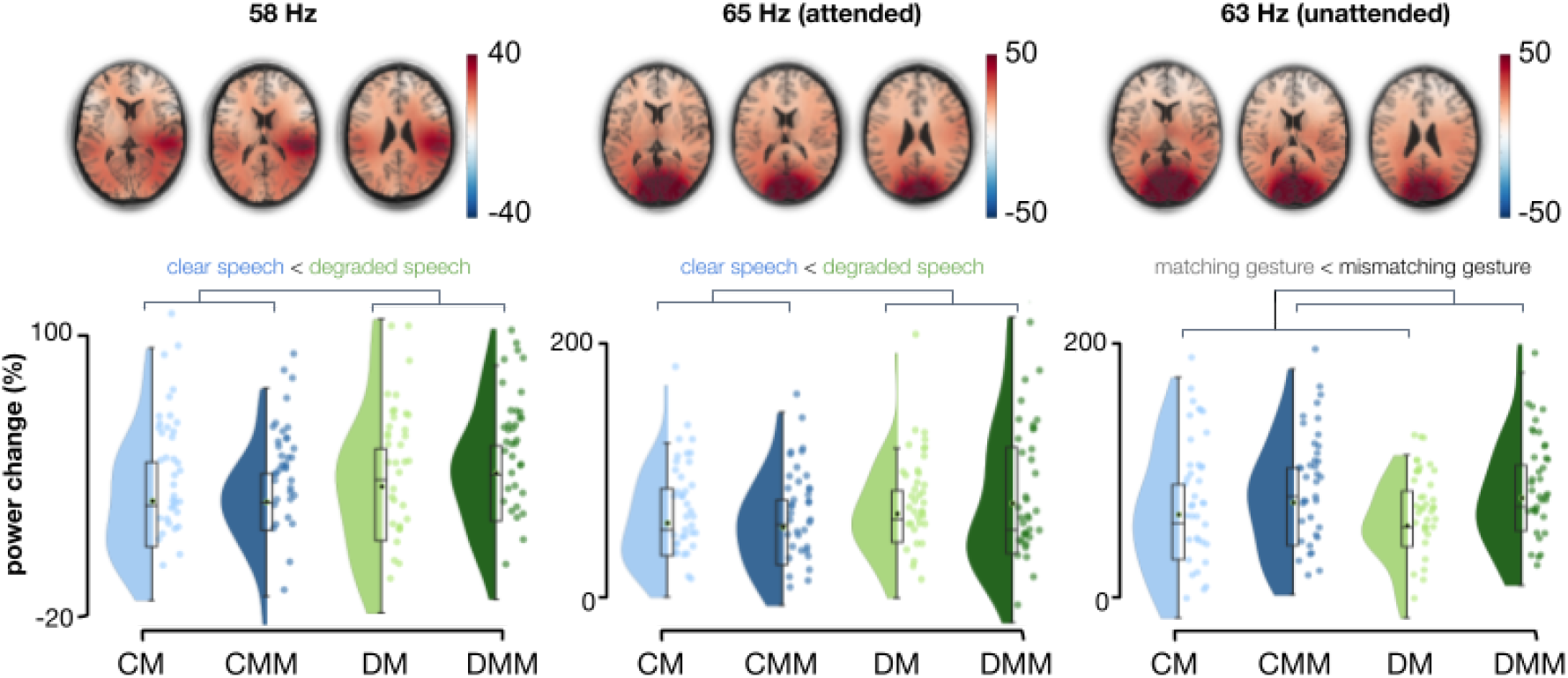
Sources of power at the auditory tagged signal at 58 Hz and the visually tagged signals at 65 Hz and 63 Hz. **A)**. Power change in percentage when comparing power values in the stimulus window to a post stimulus baseline for the different tagging frequencies, pooled over conditions. Power change is the largest over temporal regions for the auditory tagging frequency, and largest over occipital regions for the visually tagged signals. B) Power change values in percentage extracted from the ROIs. Raincloud plots reveal raw data, density, and boxplots for power change in the different conditions. CM = clear speech with a matching gesture, CMM = clear speech with a mismatching gesture, DM = degraded speech with a matching gesture and DMM = degraded speech with a mismatching gesture.

### Auditory and visual sensory regions as the neural sources of the tagging signals

Then, we proceeded to identify the neural sources of the tagged signals using beamformer source analysis. To compare conditions, we formed ROIs by selecting those grid points exceeding a threshold of 80% of peak power change (based on all conditions pooled together). Power change values per condition and per participant were compared in a 2×2 Repeated Measures ANOVA.

### Listeners engage there auditory system most when speech is degraded

For the auditory tagging frequency (58 Hz) power was strongest in right-temporal regions, and stronger when speech was degraded compared to when speech was clear (F[1, 37] = 6.91, *p* = .012, partial *η*2 = 0.157). There was no main effect of gesture (matching/mismatching; (F[1, 37] = 0.30, *p* = 0.59, partial *η*2 = 0.008) and no interaction effect (F[1, 37] = 0.44, *p* = .51, partial *η*2 = 0.012).

### Degraded speech enhances covert attention to the gestural information (65 hz)

Similarly, power at the attended visual tagging frequency (65 Hz) was stronger when speech was degraded, compared to when speech was clear (F[1, 37] = 8.45, *p* = .006, partial *η*2 = 0.186). Again, there was no main effect of gesture (matching/mismatching; (F[1, 37] = 0.27, *p* = 0.61, partial *η*2 = 0.007) and no interaction effect (F[1, 37] = 0.93, *p* = .34, partial *η*2 = 0.025).

### Mismatching gestures enhance processing of the unattended side (63 Hz)

For the unattended visual tagging frequency (63 Hz), power was stronger when gestures mismatched the speech, compared to when the gestures matched the speech (F[1, 37] = 14.80, *p* < .001, partial *η*2 = 0.286).

### 7 Hz power peak was strongest when speech was degraded and a gesture matched the speech signal

To evaluate whether intermodulation frequencies (5 and 7 Hz in our experiment) could be observed, we then calculated the power spectra at sensor- and source-level in the stimulus time window and the post-stimulus time window. Based on previous work (Drijvers et al., 2021) we focused on left frontal sensors and LIFG. Apart from a peak at 7 Hz for the DM condition, we visually did not observe clear peaks at 5Hz, nor for the other conditions at 7Hz. (Figure 5A/B). For statistical evaluation see next section.

**Figure 5.**
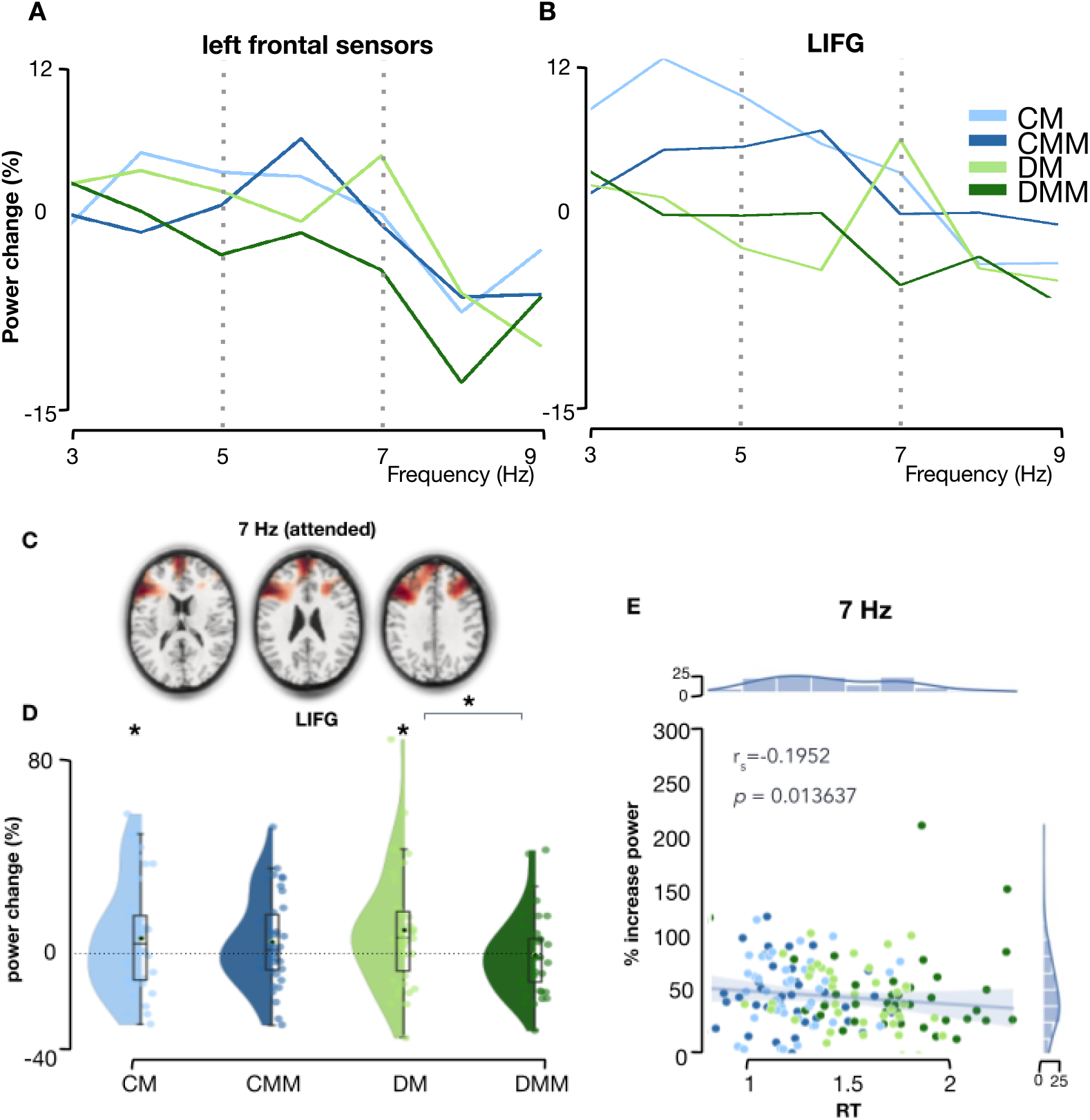
Power at the intermodulation frequencies (f_visual_-f_auditory_). **B)** Power over left frontal sensors (% increased compared to a post-stimulus baseline). B) Power over LIFG source region (% increased compared to a post-stimulus baseline) C) Sources of power at 7 Hz **D)** Power change values in percentage extracted from the Left Inferior Frontal Gyrus (LIFG) in source space. Raincloud plots reveal raw data, density, and boxplots for power change per condition. E) Correlation between power increase (% compared to baseline) and RT for each item. Colors indicate the trial type (CM, CMM, DM, DMM)

### Frontotemporal and frontal regions as the neural sources of the intermodulation signals

Beamformer source analysis confirmed left frontotemporal regions as the neural sources of the intermodulation signals. Additionally, activity in frontal regions (left/right) and in the right hemisphere was observed. To evaluate whether power at the intermodulation frequencies in LIFG was increased during the stimulus window compared to the post-stimulus baseline window, 1 sample permutation tests against zero were performed. At 7 Hz there was a significant increase in power for both conditions in which gestures matched the speech (CM: *p* = 0.044; DM: *p* = 0.009). A non-parametric Friedman test differentiated % power change across the four conditions (CM, CMM, DM, DMM), Friedman’s Q(3) = 9.126, *p* = .028. Post-hoc analyses with Wilcoxon signed-rank tests indicated increased power for the degraded match (DM) condition, compared to the degraded mismatch (DMM) condition, W = 178, *p* = .03, after Benjamini/Hochberg correction for multiple comparisons. This suggests that activity in LIFG is increased for those conditions that benefit most from integration (with a matching gesture). There were no differences between conditions at 7 Hz in the Left Postcentral Gyrus (A1/2/3), which was taken as a control region as it is not typically associated with audiovisual integration, attention, or 5-7 Hz activity related to cognitive tasks, Friedman’ s Q(3) = 0.916, p = 0.822.

### Linking the strength of the intermodulation frequency to behavior

Finally, we evaluated whether power at the intermodulation frequencies was related to the success of integration. To this end, we computed the increase in 7 Hz power in LIFG for each trial and each participant, and evaluated the Spearman’s rank correlation with the reaction times (only correct trials included). Results showed a significant negative correlation (r_s_ = -0.195, *p* = 0.014), indicating higher 7 Hz power in LIFG for trials in which participants were faster to respond (Figure 5E). These results suggest that the intermodulation frequency for the attended visual information was related to the speed of integration.

## Discussion

In the present MEG study we used RIFT to investigate how attention affects the processing of auditory and visual information, as well as their integration, during multimodal communication. Our results showed that attention selectively modulates the processing of sensory information, depending on the congruence (matching vs. mismatching gestures) and quality (clear vs. degraded speech) for the task at hand. Specifically, we observed enhanced processing of auditory information when speech was degraded. In line with previous studies (Drijvers et al., 2021) we observed a stronger drive by the 58 Hz amplitude modulation signal in auditory regions when speech was degraded compared to when speech was clear. In visual regions, we observed a stronger drive by the attended visual modulation signal (65 Hz) when speech was degraded. For the unattended visual modulation signal (63 Hz), we observed enhanced processing when gestures were mismatching. We observed enhanced activity in LIFG at the attended intermodulation frequency (7 Hz, f_visual_attended_ − f_auditory_) for those conditions that benefitted from integration (i.e. conditions with a matching gesture: CM. DM). Trials that were answered faster also showed increased activity at the attended intermodulation frequency (7 Hz) in LIFG. Together, our results suggest that attention can modulate the strength and speed of audiovisual processing and interaction, depending on the relevance and quality of the sensory input.

### Degraded speech enhances attention to auditory information

The current study provides evidence that degraded speech enhances attention to auditory information when compared to clear speech. We observed a stronger drive by the 58 Hz amplitude modulation signal in auditory cortex when speech was degraded, compared to when speech was clear. This finding is consistent with previous studies that have reported enhanced attention to degraded speech (Helfer and Freyman, 2008; Zhang et al., 2019, Drijvers et al., 2021). The increase in attention to auditory information in the degraded speech condition may be due to increased effort needed to understand the speech, leading to a greater allocation of attentional resources to the auditory signal (Wild et al., 2012).

### Degraded speech enhances processing of the attended gestural information

Additionally, degraded speech enhanced processing of the attended visual information. In occipital regions, we observed a stronger drive by the 65 Hz visual modulation signal when speech was degraded compared to when speech was clear. The enhanced attention to the attended gestural information in the degraded speech condition may be due to a compensatory mechanism, where participants rely more heavily on visual information in the presence of degraded auditory information (Drijvers & Ozyurek, 2017; Holle et al., 2010; Holle & Gunter, 2007; Ross et al., 2007; Obermeier, Dolk & Gunter, 2012; Erber, 1975; Sumby and Pollack, 1954).

In previous work, the opposite pattern was found; that is, a stronger drive when speech was clear, rather than degraded (Drijvers et al., 2021). However, in that study participants were presented with only one video in the center of the screen. This allowed for more room for participants to explore the different planes and parts of the visual information (away from the gestures). Because listeners gaze more often to the face and mouth than to gestures when speech is degraded (Drijvers, Vaitonytė, & Özyürek, 2019), this could have resulted in lower power at the visual tagging frequency when speech was degraded.

### Mismatching gestures enhance processing of the unattended gestural information

The processing of gestures during audiovisual integration has been shown to be influenced by the congruency between speech and gestural information. Our findings support this idea and suggest that the presence of mismatching gestures can reduce visual attention to the attended gestural information and enhance processing of the unattended side. This finding is consistent with previous studies showing that processing of a task-relevant stimulus can be reduced in the presence of task-irrelevant information (Lavie et al., 2004). In the current study, it is possible that the inability to integrate the mismatching gestural information led participants to allocate less attentional resources to the attended side and instead attend to the unattended side. In other words, the presence of mismatching speech gestures may reduce the allocation of attention to the attended side, potentially allowing for greater processing of the unattended side.

### Flexible allocation of neurocognitive resources

Overall, these findings suggest that the recruitment of sensory resources is not static, but dynamic. The ability to flexibly allocate neurocognitive resources allows listeners to rapidly adapt to speech processing under a wide variety of conditions (Peelle, 2018). For example, degraded speech enhances attentional allocation to both auditory and gestural information, potentially reflecting a compensatory mechanism to overcome the challenges of processing degraded speech. On the other hand, attention may be diverted when the audiovisual information does not match and therefore becomes irrelevant. These findings also highlight the importance of considering both lower-order and higher-order factors when investigating audiovisual integration and attention. The manipulation of degradation in the auditory modality allowed us to investigate the role of lower-order factors (i.e., the quality of the sensory input), while the manipulation of gesture congruence allowed us to investigate the role of higher-order factors (i.e., the semantic relationship between the auditory and visual information). Future studies could build on this by manipulating a wider range of factors such as the complexity, familiarity or timing, to better understand how different types of information interact during audiovisual processing.

### The auditory tagged speech signal and attended gestural information interact in left-frontotemporal regions

Our findings also shed light on the role of *top-down* attention in audiovisual integration. At the attended intermodulation frequency (7 Hz), we found that power in LIFG was enhanced for degraded speech with a matching gesture compared to degraded speech with a mismatching gesture. This is in line with earlier work showing an influence of the quality or relevance of sensory input in modulating audiovisual integration. For example, studies have shown that manipulations of sensory congruence can affect the degree of audiovisual integration, with greater integration occurring when stimuli are congruent across modalities (e.g., Vatakis & Spence, 2007; Welch & Warren, 1980; Talsma et al., 2010). Moreover, our results showed stronger power at 7 Hz when speech was degraded and the gesture was matching compared to when the gesture was mismatching. This suggests that when the auditory signal was weaker due to the degradation of speech, attention was shifted more strongly towards the visual modality when this was relevant, resulting in enhanced neural processing of the visual stimulus at the attended frequency. In simple audiovisual perceptual tasks, inverse effectiveness is often observed, which holds that the weaker the unimodal stimuli, or the poorer their signal-to-noise ratio, the stronger the audiovisual benefit (Kayser et al., 2005; Meredith and Stein, 1983, 1986b; Perrault et al., 2005; Stanford et al., 2005). A similar pattern has been observed for more complex audiovisual speech stimuli, where results show an enhanced benefit of adding information from visible speech to the speech signal at moderate levels of noise-vocoding (Drijvers & Ozyürek, 2017) or an enhanced benefit from bimodal presentation for words that were less easily recognized through the visual input (van de Rijt et al., 2019). In our study, in line with this idea, we observe enhanced power at the attended intermodulation frequency (7 Hz) for the degraded match condition compared to the degraded mismatch condition.

The observed effects on the intermodulation frequencies are different from earlier work (Drijvers et al., 2021) that observed a reliable peak at 7 Hz power during stimulation when integration of the lower-order auditory and visual input was optimal, that is, when speech was clear and a gesture was matching. These previous results suggested that the strength of the intermodulation frequency reflected the ease of lower-order audiovisual integration. However, results from the current study indicating an effect of gesture congruence (enhanced activity for the DM condition compared to DMM), suggest otherwise. We speculate that this discrepancy might be due to differences in task demand. Specifically, the fact that we used two videos presented on both sides of the screen, rather than a single video presented centrally as in the Drijvers et al. study, and the added attentional manipulation could have contributed to the observed differences. In the current study, because we selected specific tagging frequencies that resulted in intermodulation frequencies at 5 and 7 Hz, our effects of integration are manifested in the theta range. Theta oscillations have been implicated in both attentional selection and audiovisual integration processes. For example, theta activity seems to be related to cognitive control in cross-modal visual attention paradigms (Wang et al., 2016), multisensory divided attention (Keller et al., 2017) and theta oscillations have been shown to modulate attentional search performance (Dugue & VanRullen 2015). Thus, parts of the 7 Hz power may reflect a combination of attentional and integrative processes. For example, enhanced theta power in response to clear speech may reflect the presence of more attentional resources (driven by the simplicity of the trial). On the other hand, enhanced theta power in response to degraded speech for the attended stimulus may reflect both increased attentional demands due to the degraded speech and increased integration demands due to the need to compensate for the degraded auditory information. However, mostly mid-frontal and central brain areas, and not LIFG, have been shown to be involved in allocating and controlling the direction of attention (Yantis & Serences, 2003; Woldorff et al., 2004; Corbetta & Shulman 2002, Moore, 2003). Future studies could use different tagging frequencies (and thus different intermodulation frequencies) to try to disentangle these effects of both integration and attention.

## Conclusion

This study provides insights into the neural mechanisms underlying attentional modulation of audiovisual processing and integration during communication. By utilizing RIFT and MEG, we were able to identify the neural sources associated with sensory processing and integration, and their involvement during different requirements for audiovisual integration. Our findings highlight the critical role of degraded speech in enhancing attention to both auditory and attended gestural information, and the potential role of mismatching gestural information in shifting visual attention away from the attended side. Moreover, results suggest that the intermodulation frequency for the attended visual information was related to the speed of integration, with stronger intermodulation activity for trials with faster reaction times. Overall, our results demonstrate the complex interplay between different sensory modalities and attention during audiovisual integration and the importance of considering both lower- and higher-order factors in understanding these processes. The role of attention may be context-dependent. Understanding the factors that modulate audiovisual speech-gesture integration is crucial for developing a more comprehensive understanding of how humans communicate in daily life.

## Acknowledgements

This work was supported by a Minerva Fast Track Fellowship from the Max Planck Society awarded to LD.

## Data and code availability

Data and code to reproduce the analyses in this article are available at https://osf.io/p36xf/

